# Land productivity dynamics in and around protected areas globally from 1999 to 2013

**DOI:** 10.1101/821793

**Authors:** Begoña de la Fuente, Mélanie Weynants, Bastian Bertzky, Giacomo Delli, Andrea Mandrici, Eduardo Garcia Bendito, Grégoire Dubois

**Author notes:** corresponding author. Postal address: European Commission, Joint Research Centre (JRC), Directorate D - Sustainable Resources, Via E. Fermi 2749, I-21027 Ispra, VA, Italy.

## Abstract

Tracking changes in total biomass production or land productivity is an essential part of monitoring land transformations and long-term alterations of the health and productive capacity of land that are typically associated with land degradation. Persistent declines in land productivity impact many terrestrial ecosystem services that form the basis for sustainable livelihoods of human communities. Protected areas (PAs) are a key strategy in global efforts to conserve biodiversity and ecosystem services that are critical for human well-being, and cover about 15% of the land worldwide. Here we globally assess the trends in land productivity in PAs of at least 10 km^2^ and in their unprotected surroundings (10 km buffers) from 1999 to 2013. We quantify the percentage of the protected and unprotected land that shows stable, increasing or decreasing trends in land productivity, quantified as long-term (15 year) changes in above-ground biomass derived from satellite-based observations with a spatial resolution of 1 km. We find that 44% of the land in PAs globally has retained the productivity at stable levels from 1999 to 2013, compared to 42% of stable productivity in the unprotected land around PAs. Persistent increases in productivity are more common in the unprotected lands around PAs (32%) than within PAs (18%) globally, which may be related to more active management and vegetation cover changes in some of these unprotected lands. About 14% of the protected land and 12% of the unprotected land around PAs has experienced declines in land productivity from 1999 to 2013 globally. Oceania has the highest percentage of land with stable productivity in PAs (57%) followed by Asia (52%). Europe is the continent with the lowest percentage of land with stable productivity levels in PAs (38%) and with the largest share of protected land with increasing land productivity (32%), which may be related to the high population density and share of agricultural land within PAs as well as to rural land abandonment processes in many regions of Europe. In conclusion, we provide a relevant indicator and assessment of land productivity dynamics that contributes to characterise the state, pressures and changes in and around protected areas globally. Further research may focus on more detailed analyses to disentangle the relative contribution of specific drivers (from climate change to land use change) and their interaction with land productivity dynamics and potential land degradation in different regions of the world.

## 1. Introduction

The increasing demand of biomass for food, fodder, fibre and energy is driving changes in ecosystems globally (Vitousek et al., 1986; Erb et al., 2009; Ellis et al., 2010; Cherlet et al., 2018). The state of vegetation cover is one reliable and accepted measure associated with land productivity, since it is related to the set of ecological conditions and to the impacts of natural and anthropogenic environmental change (Cherlet et al., 2018). Therefore, changes in total biomass production or land productivity are required to characterize land transformations that are typically associated with land degradation (Yengoh et al., 2016). Persistent changes in land productivity, as captured through plant biomass production, point to long-term alteration of the health and productive capacity of land (Cherlet et al., 2018). Trends in land productivity have been adopted as one of three land-based progress indicators of the United Nations Convention to Combat Desertification (UNCCD, 2013), which are used for mandatory reporting. Land productivity trends have also been proposed as one sub-indicator for monitoring and assessing progress towards achieving the UN Sustainable Development Goal (SDG) 15, target 3 (UNCCD, 2015). Almost all terrestrial ecosystem services that sustain human livelihoods will be directly or indirectly impacted by a persistent reduction in land productivity. Declining productivity, however, is certainly not the only indicator of possible land degradation. Increased productivity can sometimes happen at the cost of other land resources, such as water or soil; in this case, it can lead to degradation that would be observable only in later stages (Cherlet et al., 2018).

Protected areas (PAs) are a key strategy in global efforts to conserve biodiversity and ecosystem services that are critical for human well-being (Watson et al., 2014; UNEP-WCMC et al., 2018). PAs are established and managed to achieve the long-term conservation of nature with associated ecosystem services and cultural values (Dudley, 2008). They play a fundamental role in the *in situ* conservation of genetic, species and ecosystem diversity, and the delivery of economic, social and cultural benefits from nature to people (Mulongoy and Gidda, 2008; UNEP-WCMC et al., 2018; Vačkář et al., 2016). The critical importance of PAs is recognized in several international agreements and targets for biodiversity conservation and sustainable development. In Aichi Target 11 of the Strategic Plan for Biodiversity 2011-2020 of the Convention on Biological Diversity (CBD), the international community agreed to conserve by 2020 at least 17% of terrestrial and inland water areas through effectively and equitably managed, ecologically representative and well-connected systems of PAs (CBD, 2010). Terrestrial PAs also contribute to the conservation target 15.1 of SDG 15 as they seek to “protect, restore and promote sustainable use of terrestrial ecosystems” (UNGA, 2015).

As of July 2018, PAs cover almost 15% of the Earth’s land area, compared to around 10% in 1992 when the CBD was signed (UNEP-WCMC et al., 2018). Thus, terrestrial PAs are globally important and are expanding their extent. Depending on their primary management objectives, PAs are commonly classified into different management categories, ranging from strict nature reserves to sustainable use areas (Dudley, 2008).

Here, we assess the trends in land productivity in PAs and in their unprotected surroundings (10 km buffers) globally from 1999 to 2013. We quantify the percentage of the global protected and unprotected land that shows stable, increasing or decreasing trends in land productivity, and examine the difference in land productivity dynamics across continents and for different PA management categories. By doing so, we provide a relevant indicator that contributes to characterise the state, pressures and changes in and around protected areas globally. We note that to identify critical land degradation zones, land productivity dynamics (hereafter LPD) must be further analysed within the context of anthropogenic land use and other environmental changes. Land productivity as here analysed and presented refers to observed changes of above-ground biomass and is conceptually different from, and not necessarily related to, agricultural production or income per unit area.

## 2. Materials and Methods

### 2.1. Protected areas and 10 km buffer zones

We downloaded the public version of the World Database on Protected Areas (WDPA) for May 2019 from Protected Planet (UNEP-WCMC and IUCN, 2019). The WDPA is managed by the World Conservation Monitoring Centre (WCMC) of the United Nations Environment Programme (UNEP) in collaboration with the International Union for Conservation of Nature (IUCN), and is collated from national and regional datasets (UNEP-WCMC, 2017). This dataset consists of 235,522 protected areas (PAs), of which those of 10 km^2^ or larger are documented in detail in the Digital Observatory for Protected Areas (DOPA) developed by the Joint Research Centre of the European Commission (Dubois et al., 2016), and are those considered in this study. The selection of this 10 km^2^ threshold is based on computational limitations in processing the large datasets involved in this study. The DOPA, accessible at http://dopa.jrc.ec.europa.eu, provides a broad range of consistent and comparable indicators on PAs at country, ecoregion and protected area level (Dubois et al., 2016). These indicators are particularly relevant for Aichi Biodiversity Target 11 (Protected Areas) of the CBD, and the UN Sustainable Development Goals 14 (Life below Water) and 15 (Life on Land).

In this study, we excluded, in addition to PAs with an area below 10 km^2^, the following PAs. First, all PAs with undefined boundaries, so-called point PAs that are reported by national authorities with only a single geographic reference for the centre of the PA. Second, we excluded all marine PAs, i.e. we only considered PAs classified in the WDPA as terrestrial and coastal (the latter comprised both terrestrial and marine portions). We only considered their land portion for all analyses. The land portion was determined using the land mask obtained from the combination of the Global Administrative Unit Layers (GAUL) for year 2015, developed by the Food and Agricultural Organization (FAO) of the United Nations^1^, and the layer on Exclusive Economic Zones (Flanders Marine Institute, 2016), according to the combination method described in Bastin et al. (2017). Third, we excluded PAs with a “proposed” or “not reported” status in the WDPA, in line with common practice for global PA analyses (e.g., UNEP-WCMC and IUCN, 2016; Saura et al., 2018). Fourth, we excluded UNESCO Man and the Biosphere Reserves, as their buffer areas and transition zones may not meet the IUCN protected area definition (Dudley 2008), and because most of their core areas overlap with other protected areas (UNEP-WCMC and IUCN, 2016). Fifth, we excluded all PAs that had not been already designated before 1999 (as well as those that had no designation year reported in the WDPA), given that the temporal period here considered is from 1999 to 2013 (see next section).

In addition, in order to compare changes within and around the PAs, we considered, around each PA, a 10 km buffer zone that did not overlap with other PAs, hereafter referred to as the unprotected 10-km buffer.

All analyses were performed by rasterizing the PAs and their unprotected buffers to a spatial resolution of 1 km, which is the resolution of the layer on land productivity dynamics, as described next.

### 2.2. Land productivity dynamics

The dataset used for the analysis of land productivity dynamics (LPD) in and around PAs is that presented in the Global Land Outlook (Sommer et al., 2017) and in the World Atlas of Desertification (Cherlet et al., 2018), which is available at https://wad.jrc.ec.europa.eu/landproductivity. This is a global map product based on vegetation phenological metrics that relate to the land’s capacity to sustain primary production. It is derived from time series of indices of vegetation photosynthetic activity, namely the Normalized Difference Vegetation Index (NDVI), obtained from satellite data acquired by the SPOT VEGETATION sensor (Ivits et al., 2013). Research has shown that time series of remotely sensed vegetation indices, such as those used for deriving LPD, are correlated with biophysically meaningful vegetation characteristics such as photosynthetic capacity and primary production. These characteristics are closely related to global land surface changes and biomass trajectories that can be associated with processes of land degradation and recovery (Yengoh et al., 2016).

The LPD map is a non-parametric combination of datasets derived from phenological metrics based on the principle of convergence of evidence. The JRC software *Phenolo* (Ivits et al.,2013a) provides the metrics for each year based on a vegetation index time series (here SPOT Vegetation NDVI 1999-2013).

The long-term change in land productivity is captured by a steadiness index (Ivits et al., 2012) combined with baseline levels and state change. The steadiness index combines the tendency of change (slope) and the net change (Multi Temporal Image Differencing (MTID) method (Guo et al. 2008)) of the growing season’s NDVI integral (*Phenolo*’s Standing Biomass) over the selected time period (here 1999-2013). The values of Standing Biomass averaged for the first three years and classified into three levels give the baseline. The state change compares the Standing Biomass at the beginning and the end of the time series (three-year average).

The recent status of the land productivity map derives from the Local Net Scaling (Prince 2004) of the land productivity. The Local Net Scaling method compares an observed value with the potential or maximal value within homogeneous land capability classes. The latter are spatial units with similar patterns of seasonal phenology and productivity dynamics that reflect both climate and land use conditions, called here Ecosystem Functional Types (EFTs) by Ivits et al. (2014). Ivits et al. (2016) stratify the global land into 100 EFTs based on a cluster analysis of phenological metrics from *Phenolo* run on NOAA GIMMS 3G NDVI time-series (1982-2010). The comparison between the potential and actual land productivity is expressed in terms of the five-year average of the potentially human appropriated vegetation production (*Phenolo*’s cyclic fraction), based on SPOT Vegetation NDVI 2009-2013.

The long term change and the current status maps are combined and reclassified into the following five classes in the LPD map, which contain information on persistent trajectories of land productivity dynamics over 15 years, from 1999 to 2013, with a spatial resolution of 1 km:

1. Persistent severe decline in productivity
2. Persistent moderate decline in productivity
3. Stable, but stressed; persistent strong inter-annual productivity variations
4. Stable productivity
5. Persistent increase in productivity

For each PA and its unprotected buffer we calculated the percentage of the land area covered by each of these five qualitative LPD classes. Areas with no photosynthetically active vegetation (i.e., hyper-arid, arctic and very-high altitude mountain regions), for which no information on land productivity trends is available, were also considered as they were part of the area covered by the PAs or their buffers. Therefore, the sum of the percentage of the area covered by each of the five productivity classes could be less than 100% because of the presence of unvegetated areas. In addition, we calculated the percentage of all land (either protected or unprotected) covered by the five LPD classes globally and in each country, as a reference for comparison of the relative prevalence of each of the classes obtained in the PAs and their buffers. To summarize the LPD results, we combined the first three classes into a single class referred to as declining land productivity for brevity.

We aggregated the LPD results, both within and around PAs, at the level of continents using the country groupings of the M49 standard of the Statistics Division of the United Nations Secretariat, available at https://unstats.un.org/unsd/methodology/m49/ (accessed May 2019). We used the classification field “region name” (here referred to as continents) in this standard, which classified the world into 6 continents. Continental values are, therefore, influenced by these country groupings, such as the Russian Federation being included in Europe (continent) and Eastern Europe (region), or Greenland being included within the Americas (continent) and Northern America (region), among other examples. We also calculated the aggregated LPD values for the European Union (EU), considering the 28 countries within the EU when this analysis was conducted (EU-28). Each PA, as well as its unprotected buffer, was considered to belong to the country reported in the ISO3 field of the WDPA. Note that the ISO3 codes from the WDPA include cases of territories under the sovereignty of other nations. Examples are Reunion Island, a French overseas territory located in the Indian Ocean, and Greenland, a self-governing territory that is part of the Kingdom of Denmark. In calculating EU values, we excluded the PAs in territories whose reported ISO3 in the WDPA was different from those 28 countries, even if under the sovereignty of a EU member state, as in the case of Greenland or Réunion Island.

We also considered, for each PA, the information available in the WDPA regarding the management category (as defined in Dudley, 2008) and summarized the LPD area values separately for each category.

## 3. Results

### Global land productivity dynamics

Globally, the extent of land with stable productivity from 1999 to 2013 is larger than that of decreasing and increasing productivity combined (Figure 1). There is more land with increasing than with decreasing productivity (Figure 1).

**Figure 1.**
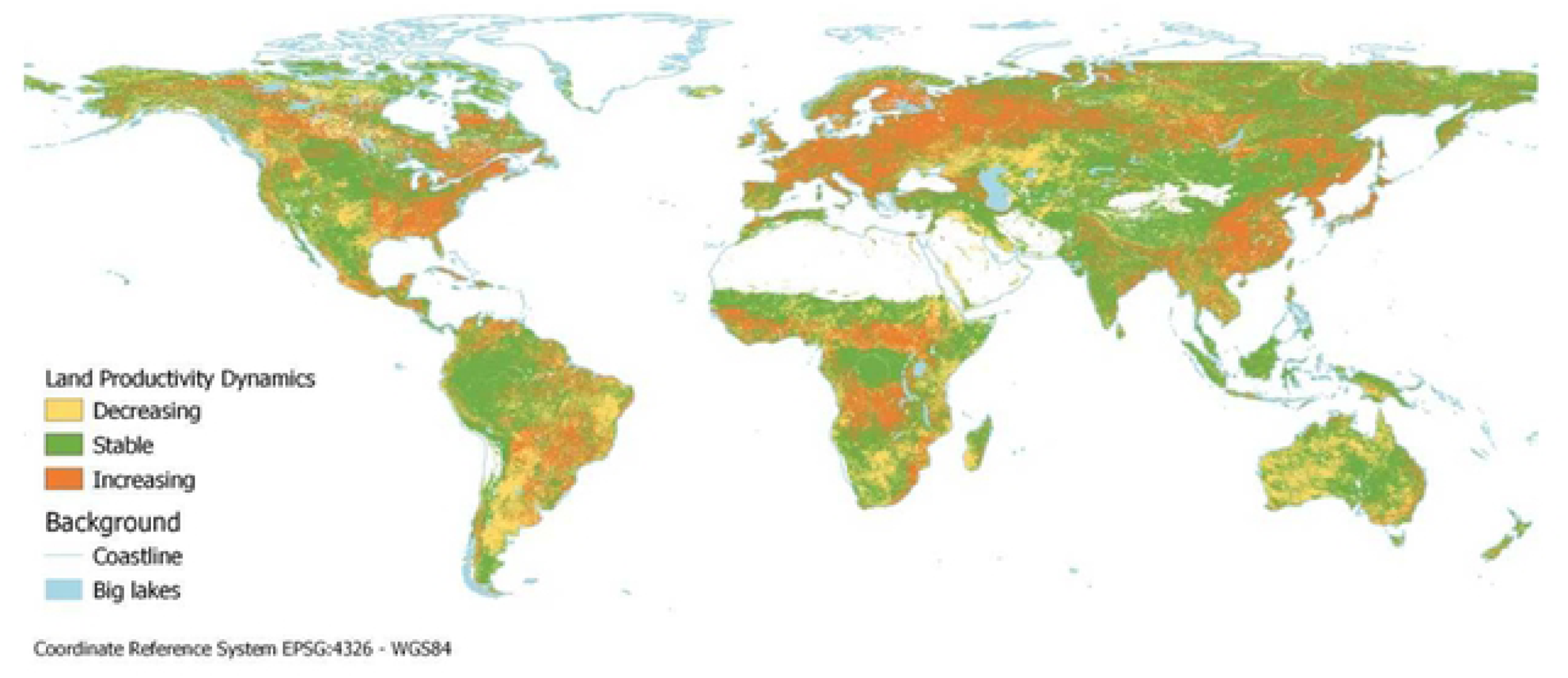
Global map of land productivity dynamics showing the spatial distribution of the three Land Productivity Dynamics (LPD) classes (decreasing – stable – increasing). The decreasing class includes areas with persistent severe decline in productivity, with persistent moderate decline in productivity and with stable but stressed productivity (persistent strong inter-annual productivity variations), as described in Methods. Land areas with no photosynthetically active vegetation are shown in white.

Global dynamics in land productivity within terrestrial and coastal PAs larger than 10 km^2^ are mainly stable (44% of the protected land), as shown in Figure 2. A similar results is found for the 10 km unprotected buffers surrounding the PAs (42%) (Figure 2). Persistent increases in productivity are more common in the unprotected land within the 10 km buffers surrounding PAs (32%) than within PAs (18%) (Figure 2). Decreasing productivity (combination of classes 1, 2 and 3 as described in Methods) is comparatively less common than stable or increased productivity: 14% of the land in PAs and 12% of the land in the unprotected buffers shows a decline in productivity (Figure 2).

**Figure 2.**
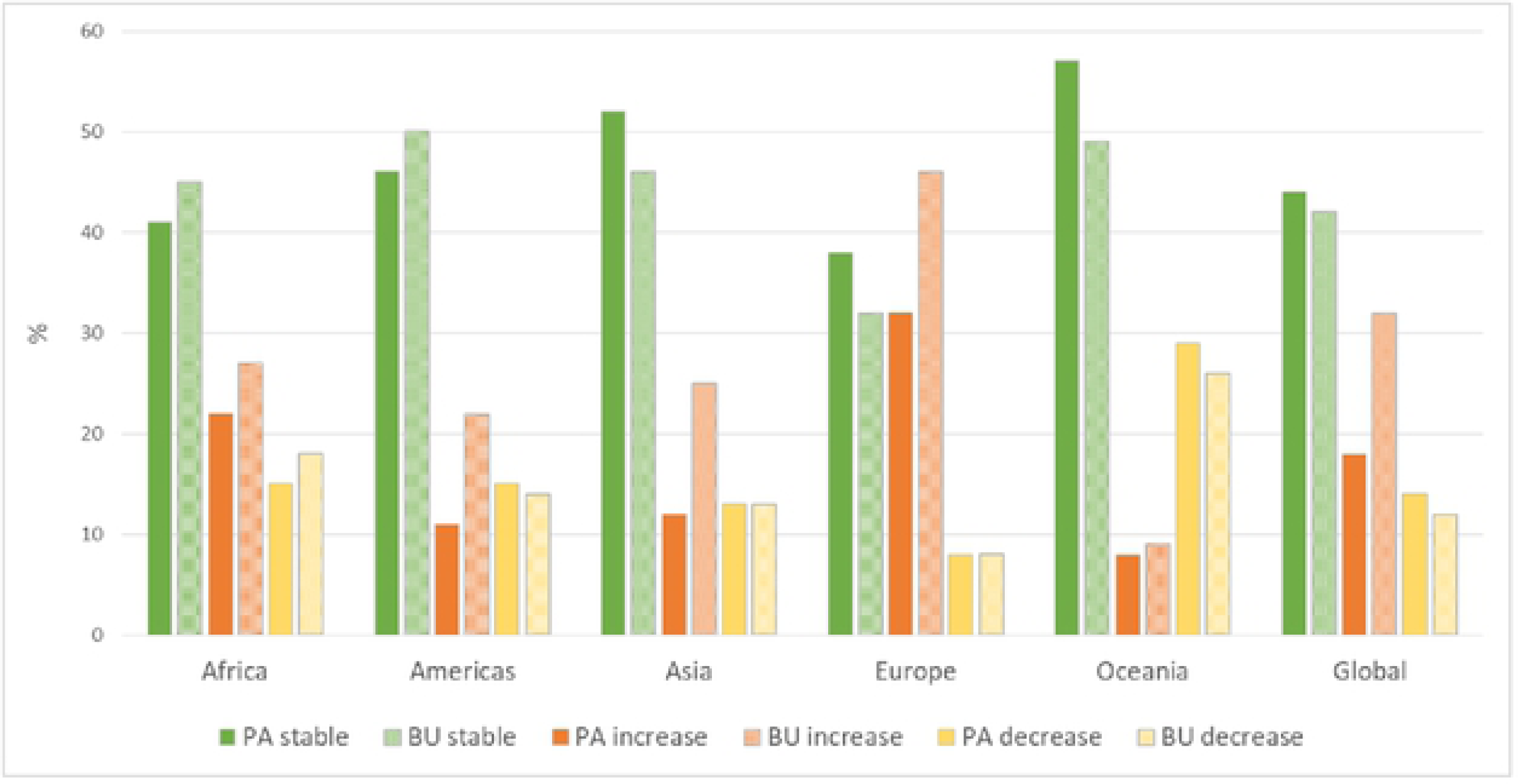
Percentage of land with stable (green), increased (orange) and decreased (yellow) land productivity between 1999 and 2013 in the protected areas (PA) and in their unprotected 10-km buffers (BU) for each continent and globally.

### Continental land productivity dynamics

The trends in land productivity within PAs are however unevenly distributed across continents. In Europe, 38% of the PA network is stable in land productivity, which decreases to 32% for the unprotected land surrounding PAs (Figure 2). These values decrease to 34% within PAs and to 30% around PAs, respectively, when focusing on the European Union (EU-28). These values are all lower than the global average; there is less land with stable productivity in and around PAs in Europe and in the EU than in any other continent (Figure 2). Africa is the continent with the second lowest percentage of land with stable productivity: 41% within PAs and 45% in the 10-km buffers around PAs (Figure 2). The highest percentage of land with stable productivity within PAs is found, at the continental level, in Oceania (57%), Asia (52%) and the Americas (46%). In all continents except Africa and the Americas, the percentage of land with stable productivity is higher within than around PAs (Figure 2).

The highest percentage of land where productivity has experienced a persistent increase in productivity from 1999 to 2013 is found within the unprotected 10 km buffer surrounding European PAs (46%), which is considerably higher than the corresponding percentage of 32% within European PAs (Figure 2). These values increase to 49% and 42%, respectively, when considering only the countries belonging to the EU. The continent with the second highest percentage of persistent increase in land productivity is Africa, with 27% of the land around PAs and 22% of the land within PAs in this change class, followed by Asia and the Americas (Figure 2). The lowest values were found in Oceania, where only 9% of the unprotected land around PAs and 8% of the land within PAs has experienced a persistent increase in productivity (Figure 2). For all continents, the percentage of land with a persistent increase in productivity is higher in the 10 km buffers around PAs than within PAs (Figure 2). Our results show a general pattern for all continents where the increase in land productivity is always higher (almost 80% higher on average) in the unprotected land surrounding PAs than within them, with a wide range of values from 13% in Oceania to 108% in Asia or 44% higher in Europe.

The percentage of land with decreasing land productivity is generally similar within and around PAs for all continents (Figure 2). Oceania is the continent where declines in land productivity are more common both within (29%) and around PAs (26%; Figure 2), while the opposite is found for Europe (Figure 2). The percentage of protected land with decreasing productivity is about three times higher in Oceania than in Europe (Figure 2).

### Global land productivity dynamics across protected area management categories

PAs with sustainable resource use (IUCN PA category VI), natural monuments (III) have the highest percentages (all over 50%) of stable land productivity within PA (Figure 3). For all IUCN categories, except wilderness areas (Ib) and national parks (II), there is more stability in the land productivity inside PAs than in the unprotected surrounding 10 km buffer (Figure 3). The largest difference in the percentage of stable land productivity between inside (61%) and outside (49%) PAs is found in category III (Figure 3), while the smallest one is found in categories V and Ia (Figure 3); for category Ia the percentage of land with stable productivity is the same inside than outside PAs. For all the categories the percentage of land with increasing productivity is higher outside PAs than within them (Figure 3). The percentage of protected land with decreasing land productivity is highest in wilderness areas (Ib,32%), while increasing land productivity is most widespread in protected landscapes (V, 31%).

**Figure 3.**
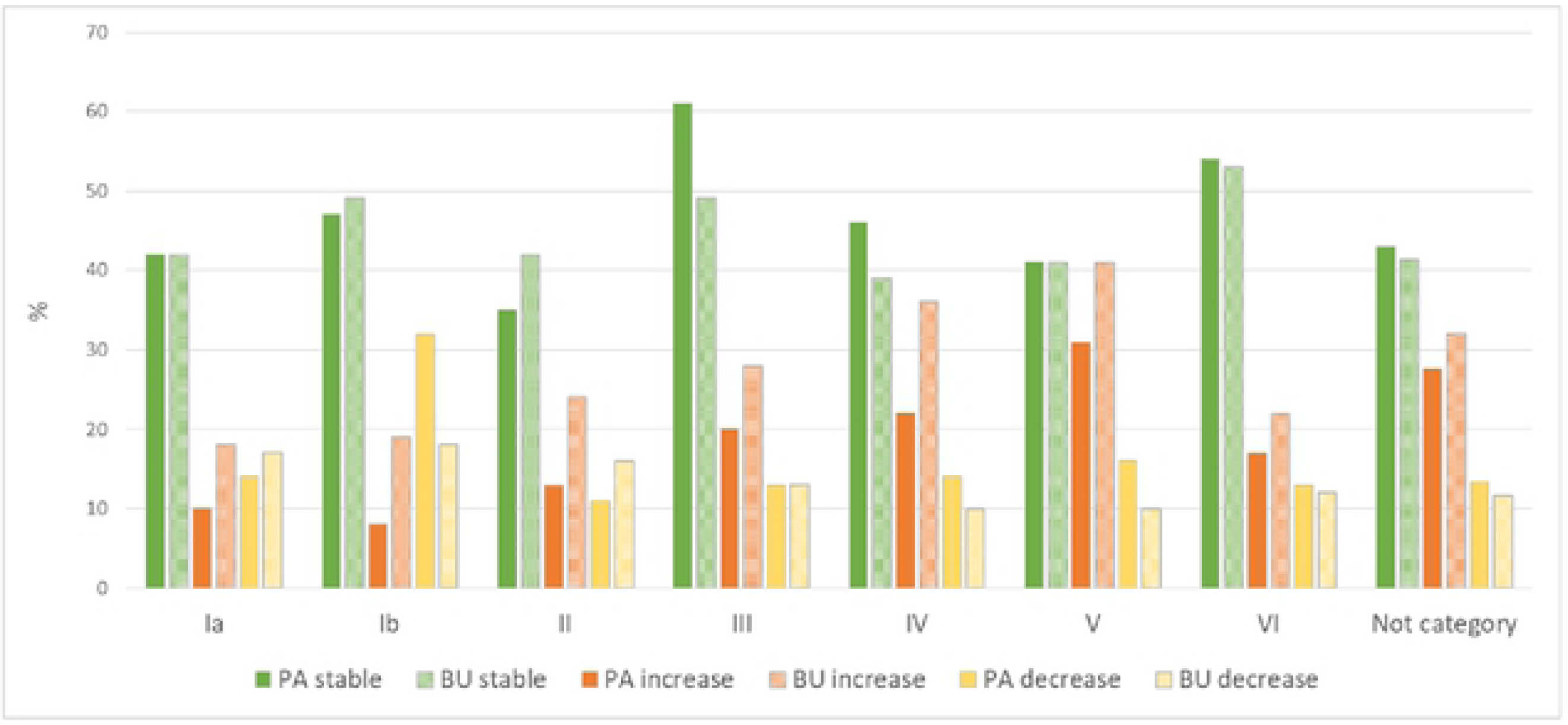
Percentage of land with stable (green), increased (orange) and decreased (yellow) land productivity between 1999 and 2013 in the protected areas (PA) and in their unprotected 10-km buffers (BU) for each IUCN management category of PAs.

## 4. Discussion

We found that 44% of the land in large (over 10 km^2^) protected areas (PAs) globally has retained the productivity at stable levels during the 15 year period here considered (from 1999 to 2013). The percentage of land with stable productivity in the 10-km buffer zones surrounding PAs only goes down to 42%. Our results, therefore, do not suggest any considerable difference between the protected and unprotected land regarding the stability in land productivity globally.

There are large tracts of land that, although being protected, have experienced declines in land productivity from 1999 to 2013. About one seventh of the protected land globally shows declines in primary productivity of above-ground biomass. This decline in land productivity, as here measured from satellite-based observations, may be related to a number of factors and processes such as deforestation, desertification or climate change, and may point to ongoing land degradation that may impact sustainability (Cherlet et al., 2018). Land degradation is however a multifaceted and complex global phenomenon with distinct variations between regions and across key land cover/land use systems which cannot be fully captured by a single indicator such as the satellite-based observed changes of above-ground biomass here considered, and would need to be explored in more detail in further research.

In the same way as declining trends in land productivity do not indicate land degradation per se, increasing trends in land productivity do not necessarily indicate a recovery or positive outcome from a conservation perspective (Cherlet et al., 2018). For instance, increased productivity is sometimes achieved at the cost of other land resources, such as water (irrigation) or soil, or is the result of the intense use of fertilizers associated with intensive agriculture (Bradford, Lauenroth & Burke, 2005). It can be also driven as a consequence of nitrogen deposition, CO_2_ enrichment fertilization, and climate change (Nemani et al., 2003; Boisvenue & Running, 2006; Thomas et al., 2010; Pettorelli et al., 2012). In several of these cases it can lead to subsequent degradation, which would be observable only in later stages. In addition, land cover and land use changes, such as decrease of primary forest cover at the expense of fast-growing tree plantations or highly-productive crops under intensive agricultural land use, can lead to an increase in the observed productivity of above-ground vegetation while having significant negative impacts on the conservation of natural resources, biodiversity and ecosystem services (Bradford et al., 2005). In rangelands and savannah, increasing productivity may be a sign of bush encroachment, which can modify the biodiversity and ecological functioning of grasslands (e.g. Leitner et al, 2018). In other ecosystems, it may arise from invasions by native or alien plant species.

For these reasons, the considerable percentage of land in PAs (18%) that has experienced an increase in productivity may be interpreted as a signal of ongoing changes that in many cases may not have benefits for the conservation objectives for which PAs have been declared. This percentage is however substantially higher in the unprotected land surrounding PAs (32%), globally almost 80% higher outside than inside. These findings may be indicative of a generally positive conservation outcome of PAs, which have avoided in many areas long-term alterations aimed to increase the productive capacity of land. The largest share of increased productivity around than inside PAs may be also indicative of more human activity or human pressure outside (Geldmann, Joppa & Burgess, 2014), related to more human settlements (De la Fuente et al., 2019) and more energy availability (Luck, 2007) that drive a more intensive use of the land. It may also be related to the fact that traditionally humans choose to settle in the most productive land (O’Neill & Abson, 2009). Other potential drivers of increase in land productivity, such as increased water supply from glacier melting or longer growing seasons in high-latitude areas driven by climate change or long-term vegetation recovery from previous natural or human-caused disturbances, may also play a significant role and may not be necessarily detrimental to biodiversity. It is however out of the reach and scope of this study to disentangle and specifically consider each of these processes, factors, and their complex interactions, which remains to be tackled in further research. In any case, the fact that the percentage of land with increased productivity is notably lower within than around PAs, as we found in this study, may be understood as a positive indicator of the relative ability of PAs to prevent changes that may pose a risk to their conservation objectives.

Interestingly, Europe is the continent with the lowest percentage of land with stable productivity levels in PAs and with the largest share of protected land with increasing land productivity. These results may be explained by the high population density and share of agricultural land use in protected areas, which is higher in Europe than in any other continent and may be associated to a significant part of the dynamics here observed (Weber & Sciubba, 2019). Also, the rural land abandonment processes have triggered the expansion of forests and woodlands in mountain areas and former agricultural lands in many European countries, often showing increasing land productivity in these areas as measured through the NDVI (Tang et al., 2011).

We emphasize that our analysis detects trends and areas with persistent declines in primary productivity that might point to ongoing land degradation, rather than areas which have already undergone degradation prior to the observation period and have reached a new equilibrium from which they do not further degrade within the observation period. Therefore, we acknowledge that our assessment may underestimate and leave unreported previous land degradation that may have occurred in PAs before 1999, which is the first year in our temporal analysis. On the other hand, the persistent land productivity changes here reported point to long-term alteration of the health and productive capacity of land. The primary productivity of a stable land system is not a steady state, but is often highly variable between different years and vegetation growth cycles due to natural variation and/or human intervention. This implies that land productivity changes cannot be assessed by comparing land productivity values of single reference years or averages of a few years. On the contrary, approaches must be based on longer term trends on multi-temporal change and trend analysis which are continuously repeated (persistent) in defined time steps using an extended time series, as is the case of the dataset and analyses used here. Despite the long-term perspective on persistent land productivity changes here adopted, we recognize that it is necessary to incorporate other factors different from biomass trends into the analysis of land degradation. To identify critical land degradation zones, land productivity must be analysed within the context of anthropogenic land use and other environmental changes.

In conclusion, we have provided a valuable assessment of land productivity dynamics in and around PAs worldwide that points to a generally positive effect of the protected area system on the conservation of land productivity. At the same time, we report that almost half of the land under protection has experienced changes (either declines or increases) in land productivity over the last 15 years. These changes may be related to a range of pressures and factors (from climate change to land use intensification) that may be detrimental for the long-term conservation of ecosystem health, biological diversity and ecosystem services. Additional and more detailed studies are needed to further analyse and disentangle the specific contribution to land productivity dynamics and potential land degradation of each of these drivers, processes and their complex interaction in different regions of the world.

## Acknowledgements

This study, the development and maintenance of the Digital Observatory for Protected Areas and the production of the land productivity map were supported mainly by the institutional activities of the Directorate D (Sustainable Resources) at the Joint Research Centre of the European Commission. We also acknowledge the contribution of the Biodiversity and Protected Areas Management (BIOPAMA) EU-ACP programme, an initiative of the African, Caribbean and Pacific (ACP) Group of States financed by the 10th and 11th European Development Funds of the European Union (EU).

## Footnotes

1 http://www.fao.org/geonetwork/srv/en/metadata.show?id=12691

